# Genetic drift drives faster-Z evolution in the salmon louse *Lepeophtheirus salmonis*

**DOI:** 10.1101/2023.12.20.572545

**Authors:** Andrew J. Mongue, Robert B. Baird

## Abstract

Sex chromosome evolution is a particularly complex sub-field of population genetics and there are still unresolved questions about how quickly and adaptively these chromosomes should evolve compared to autosomes. One key limitation to existing knowledge is an intense focus on only a handful of taxa in existing literature, resulting in uncertainty about whether observed patterns reflect general processes or are idiosyncratic to the more widely studied clades. In particular, the Z chromosomes of female heterogametic (ZW) systems tend to be quickly but not adaptively evolving in birds, while in butterflies and moths Z chromosomes tend to be evolving adaptively, but not always faster than autosomes. To understand how these two observations fit into broader evolutionary patterns, we explore, for the first time, patterns of Z chromosome evolution outside of these two well-studied clades. We utilize a publicly available high quality genome, gene expression, population, and outgroup data for the salmon louse *Lepeophtheirus salmonis*, an important aquacultural pest copepod. We find that the Z chromosome is faster evolving than the autosomes, but that this increased effect is driven by drift rather than adaptive evolution. This faster-Z effect seems to be a result of a very low effective population size of the Z chromosome, as well as high rates of female reproductive failure contributing to decreased efficiency of hemizygous selection acting on the Z. These results highlight the usefulness of organismal life history in calibrating population genetic expectations and demonstrate the usefulness of the ever-expanding wealth of modern publicly available genomic data to help resolve outstanding evolutionary questions.

## Introduction

Sex chromosomes have many fundamental characteristics that distinguish them from the rest of the genome (Abbott et al. 2017). Beyond their core function in determining an organism’s sex, they also have important roles in contributing to adaptation (Zhou and Bachtrog 2012; Mongue and Walters 2017), divergence and speciation (Presgraves 2008; Johnson and Lachance 2012; Irwin 2018), and genomic conflicts (Meiklejohn and Tao 2010; Mank et al. 2014). Moreover, they are predicted to exhibit different evolutionary dynamics relative to autosomes, which can help us to understand the forces that contribute to the evolutionary fate of genomes, as well as the consequences of this for other evolutionary processes such as adaptation and speciation.

Sex chromosomes are thought to evolve from previously homologous autosomes which acquire a sex-determining locus. Subsequent recombination suppression around the locus and degeneration of the non-recombining region leads to the formation of the Y or W chromosome (Charlesworth et al. 2005). The heterogametic sex – XY males in XY/XX systems and ZW females in ZW/ZZ systems – has only one copy of the X or Z chromosome. This has consequences for the effective population size of alleles on that chromosome, as well as the exposure of those alleles to selection. Specifically, at any given population size there are more copies of a given autosome than the X or Z in the population, meaning the latter will have a lower effective population size and will be more strongly subjected to genetic drift, and, even when recessive, they will be exposed to selection in the heterogametic sex. The combined effects of stronger genetic drift and more efficient selection are predicted to cause higher rates of nonsynonymous divergence on the sex chromosomes, i.e. a faster rate of evolution (Rice 1984; Charlesworth et al. 1987; Meisel and Connallon 2013). The so-called faster-X/faster-Z effect has been observed in a range of organisms. However, most studies are of XY systems (e.g. Ávila et al. 2014; Garrigan et al. 2014; Kousathanas et al. 2014; Charlesworth et al. 2018; Bechsgaard et al. 2019) studies on ZW systems are comparatively uncommon.

Exploration of Z chromosome evolution has focused on the two most well-studied groups – birds (Mank et al. 2007; Hayes et al. 2020; Chase et al. 2023) and Lepidoptera (moths and butterflies, e.g. Sackton et al. 2014; Pinharanda et al. 2019; Mongue et al. 2022). In birds, the faster Z effect appears to be driven by drift (Mank et al. 2009; Hayes et al. 2020; Chase et al. 2023); however in butterflies there is more argument for an adaptive faster-Z effect (Sackton et al. 2014; Mongue et al. 2022), although evidence is mixed (Rousselle et al. 2016; Pinharanda et al. 2019). Moreover, in each of these lineages the Z chromosome is conserved, i.e. all birds share a single-origin Z (Griffiths et al. 1998) and all Lepidoptera share a different orthologous Z (Fraïsse et al. 2017). In other words, there is no consensus on how Z chromosomes evolve and the contradictory evidence amounts to a disagreement between a single pair of evolutionarily independent observations. To better understand Z chromosome evolution requires investigation of a broader range of taxa.

Crustacea are an ancient (∼500mya, Zhang et al. 2007), polyphyletic clade (Regier and Shultz 1997) with diverse sex determination systems (Ye et al. 2023). Most known categories of sex determination system are represented among the Crustacea, including male (Becking et al. 2017) and female (Jiang and Qiu 2013) heterogamety, sequential (Chiba 2007) and simultaneous hermaphroditism (Hessler et al. 1995), environmental sex determination (Kato et al. 2011), polygenic sex determination (Voordouw et al. 2005; Richardson et al. 2023), asexual reproduction (Boyer et al. 2023), parthenogenesis (Elkrewi et al. 2022), as well as control by non-Mendelian genetic elements like *Wolbachia* (cytoplasmic sex determination, (Cordaux et al. 2011). ZW sex determination occurs in three major crustacean clades: Malacostraca (e.g. in isopods, (Juchault and Rigaud 1995), Branchiopoda (e.g. in fairy shrimp, (Huylmans et al. 2019), and Maxillopoda (e.g. copepods, (Borchel et al. 2022). In all three cases, ZW sex determination may have evolved independently, since each group also contains members with alternative sex determination mechanisms (Ye et al. 2023). Moreover, Maxillopoda and Malacostraca are distantly related from Branchiopoda, with phylogenetic studies supporting the grouping of the latter clade with other Arthropoda, including insects (Regier et al. 2010; Giribet and Edgecombe 2019). Thus, crustaceans offer ample opportunities to test predictions of sex chromosome evolution in diverse and ancient clades where sex chromosomes have likely independent origins; the only limitation is investment in genomic and transcriptomic resources to uncover patterns of molecular evolution.

Aside from laboratory models, the best studied and most sequenced taxa tend to be those that have some economic relevance to humans, including aquacultural pests. Some copepods are parasitic fish pathogens that spend the majority of their life cycle on their hosts. The salmon louse *Lepeophtheirus salmonis* is a parasite of exclusively salmonid hosts and is currently thought to be one of the most important hindrances to salmon aquaculture (Taranger et al. 2015; Myksvoll et al. 2018), for instance causing an estimated annual loss of >£300 million to the salmon farming industry in Norway alone (Brooker et al. 2018).

Using resources generated to better understand and control this pest copepod, we present a study of faster-Z evolution in the copepod salmon louse *Lepeophtheirus salmonis* with an aim to broaden understanding of molecular evolution of sex chromosomes. We find evidence of faster-Z evolution which appears to be driven by increased drift on the sex chromosome rather than increased adaptation. To our knowledge, this is the first study of faster-Z evolution outside of birds and Lepidoptera and demonstrates how an increasing wealth of genomic data, generated for disparate reasons, can be used to answer outstanding questions in evolutionary genetics.

## Materials and methods

### Data used

We used a previously-generated *L. salmonis* reference genome, corresponding annotation and male and female Illumina WGS DNA resequencing data (Joshi et al. 2022, see **Table 1** for accessions), as well as publicly-available male and female Illumina RNAseq libraries from whole body, antennae and gonadal tissue (Heggland et al. 2020; Skern-Mauritzen et al. 2021). For outgroups, we used publicly available WGS HiFi data for *L. nordmanni*, and also generated WGS data from a single *L. parviventris* female individual. To this end, genomic DNA was extracted using an Omniprep Kit (G Biosciences) and sequenced with Novogene Inc. (Sacramento CA, USA) for 150bp paired-end Illumina reads. All Illumina FASTQ files were trimmed using fastp v0.2.1 (Chen et al. 2018) prior to downstream analyses.

**Table 1.** Summary of data used in analyses. All data for this project are publicly available at the accessions listed above. All except for the *L. parviventris* sequencing (which was ultimately not used in analyses) were previously available before this project, demonstrating the opportunities for evolutionary genetic research using existing datasets.

| Data component | Description | Accession | Citation |
| --- | --- | --- | --- |
| Genome assembly | Chromosomal with unplaced smaller scaffolds | GCA_905330665.1 | (Joshi et al. 2022) |
| Gene annotation |  | GCA_905330665.1 |  |
| DNA resequencing | Male for polymorphism | SRR6913717 |  |
|  | Male for polymorphism | SRR6913718 |  |
|  | Male for polymorphism | SRR6913719 |  |
|  | Male for polymorphism | SRR6913720 |  |
|  | Female to validate Z linkage | SRR6913715 |  |
| Outgroup sequence | Pacbio reads for *Lepeoptheirus nordmanii* | ERS5277536 | Genome note forthcoming |
|  | Illumina reads for *Lepeoptheirus parviventris* | Available upon publication | This article |
| Male RNAseq | Adult, whole bodies | SRX10236665 | (Heggland et al. 2020; Skern-Mauritzen et al. 2021) |
|  | Adult, antennae | SRX10230935 |  |
|  | Adult, testis | SRX10230933 |  |
|  | Preadult, whole body | SRX7025567 |  |
|  | Preadult, whole body | SRX7025566 |  |
|  | Preadult, whole body | SRX7025565 |  |
| Female RNAseq | Adult, whole body | SRX4605685 |  |
|  | Adult, antennae | SRX10230937 |  |
|  | Adult, ovaries | SRX10230932 |  |
|  | Preadult, whole bodies | SRX3290140 |  |
|  | Preadult, whole bodies | SRX3290139 |  |
|  | Preadult, whole bodies | SRX3290138 |  |

### Verification of sex linkage

Because the genome assembly was not entirely assigned to chromosomes, we tested unassigned scaffolds for potential Z linkage. We aligned one male and one female DNA resequencing sample to the reference genome independently using Bowtie2 v2.4.1 (Langmead and Salzberg 2012). We used BEDtools v2.4.1 (Quinlan and Hall 2010) to calculate mean per-base coverage in 100 kilobase windows for these alignments. We then calculated the difference in coverage between the sexes as the log_2_(male/female) coverage ratio with the logic that autosomal scaffolds should have roughly equal coverage (log_2_(1) = 0) whereas the Z chromosome, which is diploid in males and haploid in females should be male-biased in coverage (log_2_(2/1) = 1). Because coverage can be variable on short scaffolds or misassemblies which join partly autosomal and partly Z-linked sequences, we introduced the criteria that the standard deviation in coverage across the scaffold should be < 0.1 to avoid sex-linkage assignment on noisy scaffolds. Nevertheless, these short scaffolds should also carry relatively few genes, making their placement relatively unimportant.

### Differential gene expression analysis

We chose a set of publicly available RNAseq samples to be comparable between the sexes (**Table 1**). Specifically, we used adult whole bodies, adult ovaries and testes, adult antennae, and a set of three preadult whole body RNA samples for each sex. While these samples are biologically comparable between the sexes, a number of idiosyncrasies prevented us from using them for traditional differential expression analyses. Most importantly, only the preadult samples have the requisite biological replicates; however, sex-biased gene expression tends to be understated in juveniles compared to adults, making the unreplicated adult tissues indispensable. Additionally, even within the replicated data, male samples were single individuals while female samples were apparently pooled, creating an uneven level of replication between the sexes.

To make the most of these data, we first quantified expression against the annotated gene set (Joshi et al. 2022) using Kallisto (Bray et al. 2016). We then used raw counts of transcripts to categorize genes as sex-biased on the basis of the SPM metric (Kryuchkova-Mostacci and Robinson-Rechavi 2017) in R (R Core Team 2023). In brief, this metric, the square of a given gene’s expression value in the focal tissues (e.g. female RNAseq) divided by the square of the expression in all tissues, quantifies genes on a scale from 0 to 1. In the example of female tissue as the focus, an SPM of 1 indicates 100% female-specific expression, whereas SPM of 0 would be 100% male-specific. We defined genes with SPM > 0.8 in females as female-biased and those with SPM < 0.2 in females as male-biased; genes in-between we categorized as unbiased. This SPM metric has proven robust and comparable to traditional DEseq analysis in previous sex chromosome work (Mongue et al. 2022).

### Analysis of sex chromosome dosage effects

Some explanations for sex chromosome evolution rely on how ploidy (Z females vs ZZ males) affects overall levels of gene expression on the sex chromosomes. Because Z chromosomes are unstudied in sea lice, if the Z chromosome shows a dosage effect between the sexes. To explore this pattern, we used the same set of male and female RNAseq samples described above, but this time averaged FPKM (fragments per kilobase of transcript per million mapped reads) across all samples for a given sex. Next, because our goal was to compare overall mean and median expression between the sexes, we excluded genes that had zero average FPKM in either males, females, or both sexes. Finally we used a Mann-Whitney-Wilcoxon test to compare expression levels between the Z chromosome and autosomes within each sex.

### Sex-bias composition of the Z chromosome

Many of the predictions for sex chromosome evolution and adaptation are predicated on the sex chromosomes having different compositions of sex-biased genes (i.e. those expressed mostly or entirely in one sex) than the autosomes. To explore the composition of the sex chromosomes, we used the sex-bias assignments from our differential expression analysis above and performed a simple Chi-squared test of independence to test if the Z chromosome differed from the autosomes as a whole in the proportion of sex-biased genes.

### Population genomic analysis

To analyze divergence (i.e. *dN/dS* ratios), needed a closely related congeneric species for comparison, but the phylogenetic relationships and divergence rates are poorly characterized in sea lice. To maximize our chance of useful inference, we tried to estimate divergence with two other *Lepeophtheirus* species, *L. nordmanii* and *L. parviventris.* The former was sequenced for the Darwin Tree of Life Project and had publicly available HiFi reads (**Table 1**) which we aligned to the reference with Minimap v2.17-r941 (Li 2018). The latter we obtained from a museum collection at the University of Florida’s Florida Museum Marine Invertebrate Collection. These specimens were originally collected by Gustav Paulay from San Juan Island in Washington State, USA in October of 2011. We aligned these data to the *L. salmonis* reference, but did not proceed further based on low alignment rates (<10%), suggesting *L. parviventris* was too diverged to be useful in this study.

After deciding to proceed with *L. nordmanii*, we processed alignment files (sorted, duplicates marked and removed, reads groups added) using Picardtools (http://broadinstitute.github.io/picard/) and called and genotyped variants with GATK-4 (Van der Auwera and O’Connor 2020), using the best-practices pipeline (McKenna et al. 2010; DePristo et al. 2011), resulting in 1,483,577 single-nucleotide variant (SNV) calls. We filtered these (Quality by Depth >2, Fisher Strand-bias <60, Mapping Quality >40) with GATK-4 SelectVariants, resulting in 1,259,828 SNVs, and then annotated and variants using SnpEff (Cingolani et al. 2012b) and extracted variant information with SnpSift (Cingolani et al. 2012a). We used a custom R script (Mongue et al. 2022) to annotate codon degeneracy for all genic sites in order to weigh variant counts by their degeneracy. Finally, we used a custom R script to calculate the ratio between non-synonymous variants per non-synonymous site (*dN*) and synonymous variants per synonymous site (*dS*). The same process was applied to the *L. salmonis* resequencing data to obtain counts of non-synonymous polymorphisms per non-synonymous site (*pN*) and synonymous polymorphisms per synonymous site (*pS*). Finally, α (alpha), i.e. the proportion of substitutions driven by adaptive evolution, was calculated as (*dN/dS*) / (*pN/pS*) (Eyre-Walker et al. 2006).

To test for significant differences between population genetic statistics, we used non-parametric tests, Mann-Whitney-Wilcoxon for pairwise comparisons and Kruskal-Wallis tests for comparisons between more than two groups. In the event of a significant Kruskal-Wallis test, we performed a post-hoc Dunn test to assess which categories differed in a pairwise framework. Finally, to assess differences in α, we used non-parametric bootstrapping and permutation tests to assess how rare differences in point estimates of α should be based on random assignment of genes into different classes (e.g. autosomal vs Z-linked). This method is based on previous population genetic investigations (Mongue et al. 2019, 2022).

We also estimated the effective population size of the Z relative to the autosomes (i.e. *Ne*_Z_/*Ne*_A_) To this end, we identified all fourfold degenerate (i.e. putatively neutral) sites and estimated heterozygosity (Watterson’s Θ) at these sites using the resequencing alignments and the population genomics tool ANGSD (Korneliussen et al. 2014). We calculated *Ne*_Z_/*Ne*_A_ as mean Θ across fourfold degenerate sites on the Z contigs divided by mean Θ across the fourfold degenerate sites on the core autosomal contigs.

## Results

### Updates to sex linkage of unplaced scaffolds

We used male and female resequencing data to validate the Z-linkage in the reported assembly as well as assign any non-chromosomal scaffolds that may have been overlooked initially. We recovered the main Z scaffold, HG994594.1, as well as a handful of other smaller scaffolds. We considered a scaffold as Z-linked if it had a log2 M/F coverage ratio of >0.9 and a coverage standard deviation < 0.1. These criteria identified CAJNVT010000004.1, CAJNVT010000006.1, CAJNVT010000008.1, CAJNVT010000010.1, and CAJNVT010000011.1 as Z-linked. These smaller scaffolds have only 21 genes on them in total but still represent a marginal improvement in the accuracy of sex linkage. In total, we assigned 23.7 Mb as Z-linked and the remaining 608.5 Mb as autosomal.

### The Z chromosome is balanced but uncompensated

We compared the average FPKM of autosomal and Z-linked genes within males and females separately. In both cases, the Z chromosome was more lowly expressed than the autosomes (W = 3101777, p = 0.018 for females; W = 3132811, p = 0.0064 for males). Directly comparing expression between the sexes on the Z, we found no difference between males and females (W = 77631, p = 0.254). In other words, the dosage of the Z chromosome is balanced between the sexes, but not compensated relative to autosomal levels.

### The salmon louse Z is more sex-biased than the autosomes

Based on our SPM metrics, we identified 1,716 female-biased, 3,439 male-biased, and 10,016 unbiased genes. These genes are not randomly distributed throughout the genome; the Z chromosome holds significantly more sex-biased genes than the autosomes (□^2^_2_ = 20.02, p = 0.00005). In particular, the Z holds more male-biased genes and marginally more female-biased genes than the autosomes. Consequently, there are proportionally fewer unbiased genes on the Z than the autosomes (**Table 2**). Taken to the extreme, we recovered 244 genes on the autosomes with female-limited expression, and a further 8 on the Z chromosome. In comparison, we found 478 autosomal male-limited genes with 19 on the Z chromosome.

**Table 2.** Distribution of sex-biased genes in the L. salmonis genome. Compared to the autosomes, the Z has significantly more sex-biased genes and fewer unbiased genes. This pattern is driven primarily by a wealth of male-biased genes, but there are marginally more female-biased genes, proportionally, on the Z as well.

|  | Female-biased | Unbiased | Male-biased |
| --- | --- | --- | --- |
| Autosomes | 1656 (11.2%) | 9788 (66.3%) | 3318 (22.5%) |
| Z chromosome | 60 (14.8%) | 223 (55.2%) | 121 (30%) |

### The salmon louse Z is faster evolving between species than the autosomes are

We recovered a faster Z effect in *L. salmonis* based on a Mann-Whitney-Wilcoxon test test of the scaled rate of divergence, *dN*/*dS* between the Z and the autosomes (W = 587246, p = 0.0015; **Figure 1**, left). When considering sex-biased genes separately, we found that female-biased, unbiased, and male-biased genes do not evolve at different rates on the Z (Kruskal- Wallis □^2^_2_ = 0.017, p = 0.991, **Figure 1**, right top). On the autosomes however, we did recover a difference (Kruskal-Wallis □^2^ = 25.40, p = 0.000003), with female-biased genes evolving faster than male-biased genes (p < 0.00001) or unbiased genes (p < 0.00001), but unbiased and male-biased genes not distinct from each other (p = 0.214, **Figure 1**, right bottom).

**Figure 1.**
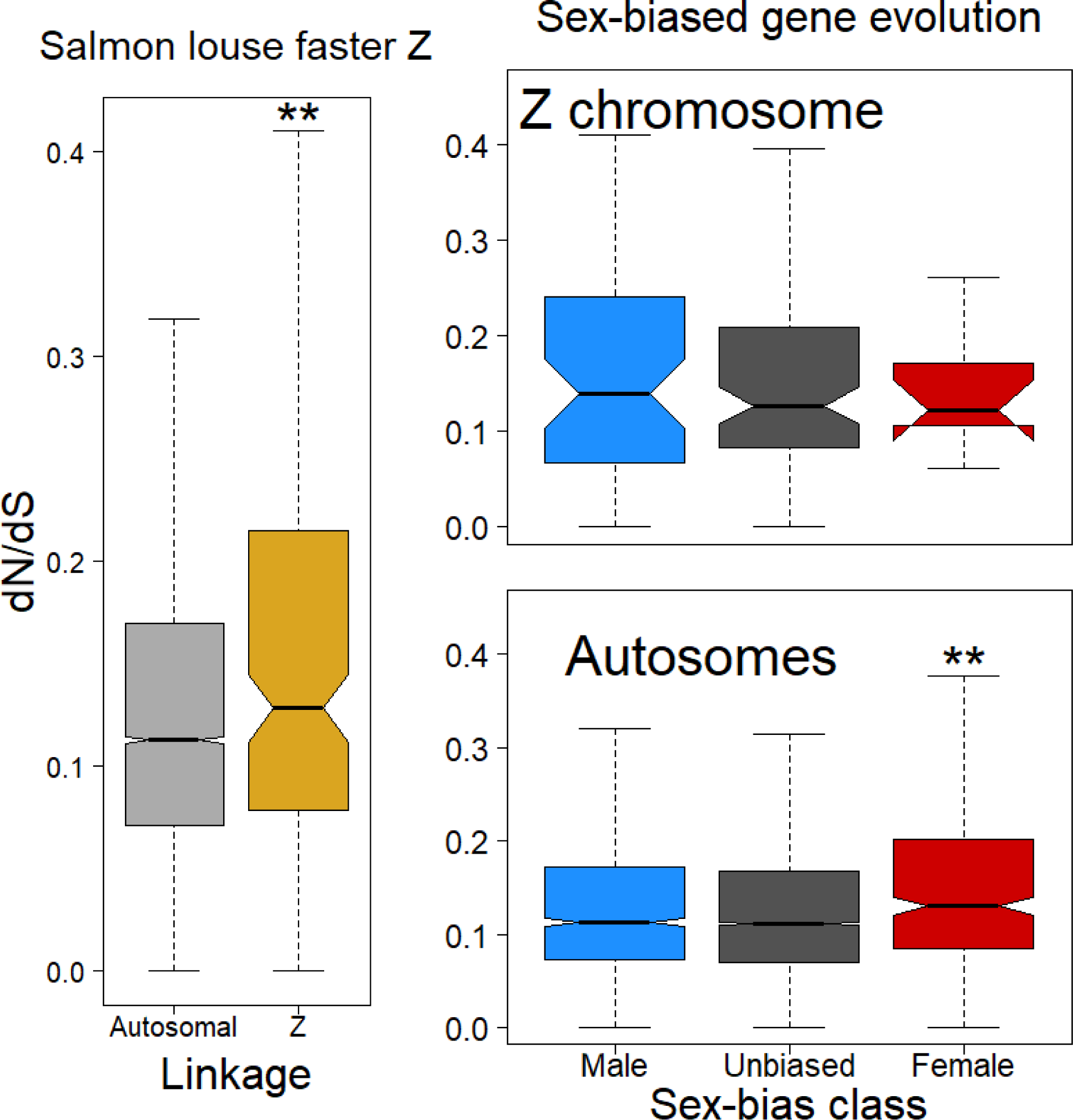
Long-term evolutionary dynamics in *Lepeophtheirus*. Left: The Z chromosome evolves more quickly (has higher *dN*/*dS*) than the autosomes overall. **Right top:** Genes of differing sex-biased expression do not evolve differently on the Z however. **RIght bottom:** But on the autosomes, female-biased genes evolve more quickly than male-biased or unbiased genes.

### The salmon louse Z holds comparable rates of scaled polymorphism to the autosomes

We next explored the same patterns using within-species variation, i.e. polymorphism data. We found a marginal but not strong trend for increased scaled polymorphism (*pN*/*pS*) on the Z compared to the autosomes (p = 0.0653, **Figure 2**, left). Given the trend and in order to more easily make comparisons with divergence data, we next explored rates of polymorphism of sex-biased genes on the Z and autosomes separately. We found that the sex-biased genes evolve differently from each other on both the Z (Kruskal-Wallis □^2^_2_ = 7.02, p = 0.0299) and autosomes (p < 0.000001). On the Z chromosome, this result is driven by greater scaled polymorphism in female-biased genes compared to male-biased (p = 0.0083) or unbiased genes (p = 0.0329); male- and unbiased genes do not hold significantly different amounts of variation (p = 0.328, **Figure 2**, right top). On the autosomes, all three classes hold different amounts of variation, with female-biased having the most (p <0.00001 vs both male-biased and unbiased), followed by male-biased genes (p < 0.00001 vs. unbiased genes), and with unbiased genes having the lowest *pN*/*pS* (**Figure 2**, right bottom).

**Figure 2.**
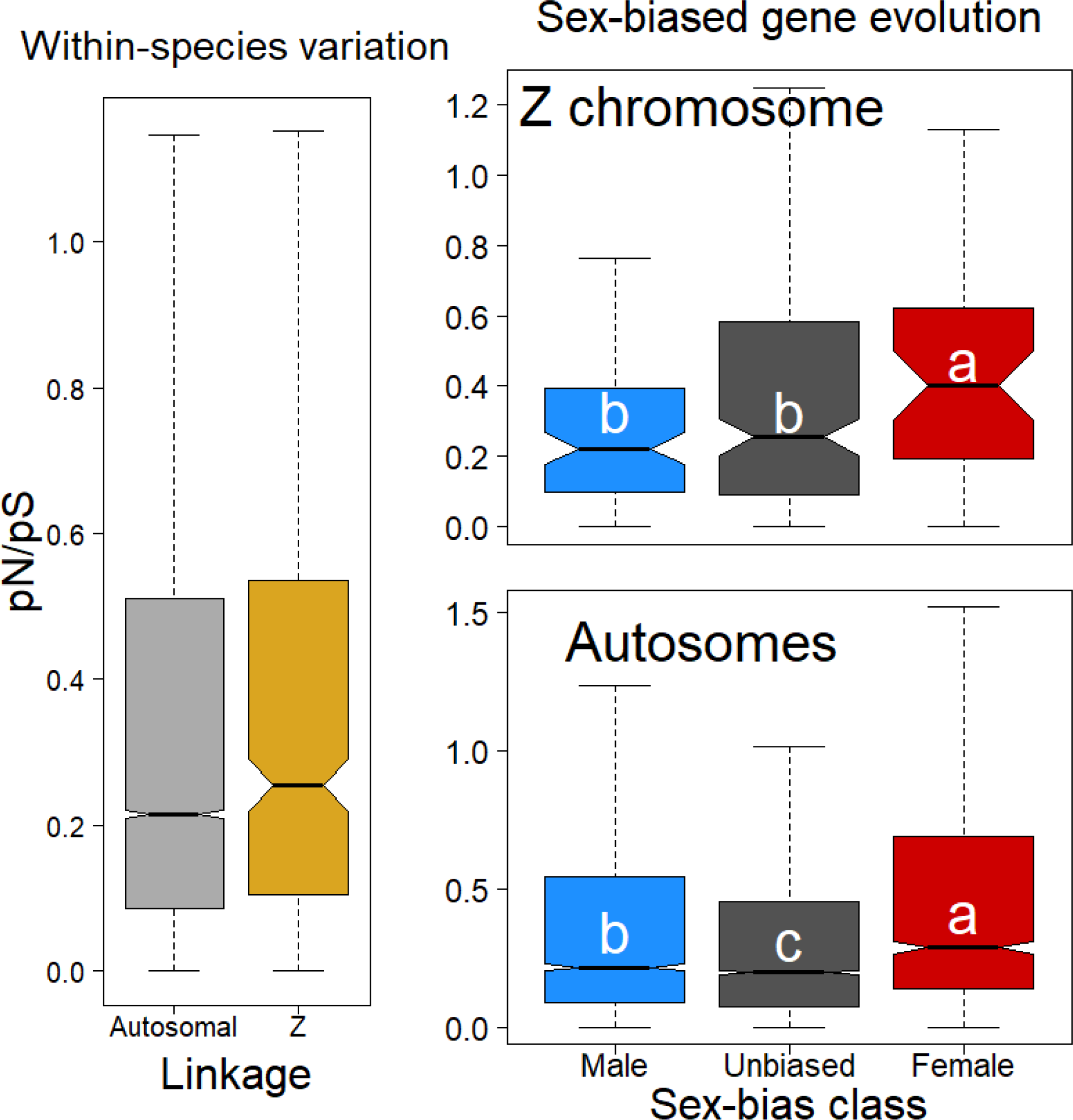
Short-term variation in *Lepeophtheirus salmonis*. Left: The Z chromosome holds comparable, but marginally more scaled non-synonymous variation compared to the autosomes. **Right, top:** Female-biased genes on the Z chromosome have higher *pN*/*pS* than unbiased or male-biased genes. **Right, bottom:** On the autosomes, female-biased genes similarly have the highest scaled non-synonymous polymorphism, followed by male-biased and then unbiased genes. Letters indicate significant differences between groups and follow the convention a > b > c.

### The faster Z is not driven by increased adaptation

By relating long-term (*dN*/*dS*) variation with short-term (*pN*/*pS*) variation, we tested whether the faster Z effect is driven by increased selection on Z-linked genes. Overall, there was no difference between the proportion of substitutions driven by positive selection, α , between the Z and autosomes (p = 0.997 by permutation). Given the lack of overall difference, we focused on comparisons between sex-bias classes within a linkage category, i.e. within autosomes and within the Z (**Table 3**). We note that our reported α values are negative, which is not defined under the strict definition of the statistic, i.e. as a proportion of adaptive substitutions its value cannot be negative. Negative α values are often ascribed to an excess of weakly deleterious segregating variants that will never fix in the population (Messer and Petrov 2013), and though these values create a downward bias in the statistic, they nevertheless represent real-world patterns of variation. Moreover, they do not invalidate within-genome comparisons between α values (Mongue et al. 2019). In other words, because we are more interested in making relative comparisons about adaptive evolution than in accurately estimating the true proportion of adaptive amino acid substitutions, negative α values are of little concern.

**Table 3.**
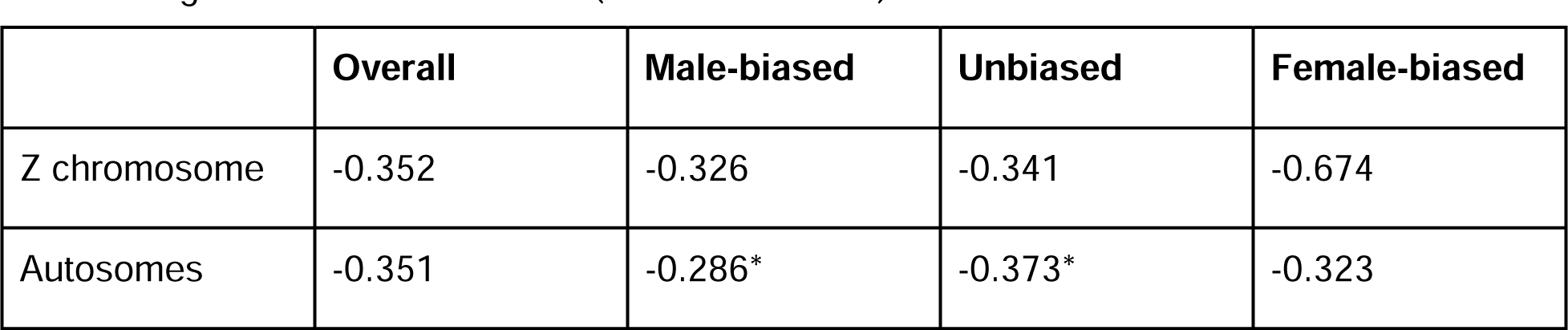
Point-estimates of adaptive evolution, α. Estimates were highly consistent across the genome and across sex-bias classes with the only significant difference being male vs unbiased genes on the autosomes (marked with an *).

|  | Overall | Male-biased | Unbiased | Female-biased |
| --- | --- | --- | --- | --- |
| Z chromosome | -0.352 | -0.326 | -0.341 | -0.674 |
| Autosomes | -0.351 | -0.286* | -0.373* | -0.323 |

On the Z, male-biased genes do not show different signatures of adaptation than unbiased genes (p = 0.950) or female-biased genes (p = 0.412). Unbiased genes and female-biased genes also did not evolve differently (p = 0.458). On the autosomes, we found male-biased genes evolve more adaptively than unbiased genes (p = 0.019), but no differently than female-biased genes (p = 0.539). Female-biased genes likewise did not evolve differently than unbiased genes (p = 0.437).

### Effective population size of the Z chromosome

Under null assumptions, the effective population size of the Z relative to the autosomes (i.e. *Ne*_Z_/*Ne*_A_) should equal ¾ (Sayres 2018). However, observed *Ne*_Z_/*Ne*_A_ may vary considerably, for example as a result of variance in reproductive success (Vicoso and Charlesworth 2009) or recombination rates between sex chromosomes and autosomes (Langley et al. 1988). We estimated *Ne*_Z_/*Ne*_A_ in *L. salmonis* as 0.95, higher than expected under naïve predictions.

## Discussion

Aligning theoretical predictions with biological observations has remained challenging for the evolution of sex chromosomes (Meisel and Connallon 2013; Charlesworth et al. 2018). The true extent to which theory and data disagree is somewhat unclear for a number of confounding reasons including methodological differences in analyses and different resolutions of datasets (e.g. much of the disagreement of faster-Z in Lepidoptera comes from a study that did not use whole-genome data; Rousselle et al. 2016). More fundamentally, many independent studies focus on just a handful of sex chromosome systems with shared evolutionary origins, such as the Z in birds (Hayes et al. 2020; Chase et al. 2023) or Lepidoptera (Sackton et al. 2014; Mongue et al. 2022), or the X in *Drosophila* (Ávila et al. 2014; Garrigan et al. 2014), making observations evolutionarily non-independent. To help address this knowledge gap, we have explored the evolutionary dynamics of the sex chromosomes of a previously unstudied clade with an independently evolved Z chromosome, the salmon louse within Copepoda.

To begin with, our results align well with common predictions for sex chromosomes: namely that they should be faster evolving and harbor more sex-biased genes than the autosomes. While perhaps not surprising, it is worth noting that these results are not universally recovered for either the speed (Rousselle et al. 2016; Whittle et al. 2020; Mrnjavac et al. 2023) or composition (Vicoso and Charlesworth 2006) of the sex chromosome. Adding more evidence from a previously unexplored lineage adds robustness to the pattern of a faster and more masculinized Z chromosome. Traditional theory has emphasized that the direction of sex-bias on the sex chromosomes should depend on the dominance or recessivity of sex-biased alleles; for instance the Z should accumulate dominant male-biased genes but also recessive female-biased genes (Rice 1984; Klein et al. 2021). While logically sound, predicting sex-biased composition of the sex chromosomes has been difficult *a priori* because the distribution of dominance effects of alleles is always unknown. Our results alongside many others suggest that many male-biased alleles are in fact dominant or at least additive, given that the Z is masculinized in almost every known case (Storchová and Divina 2006; Ellegren 2011; Mongue and Walters 2017).

### A non-adaptive Z in salmon lice

There is considerably less agreement on the role of adaptation in Z chromosome evolution, however. To a first approximation, the Z chromosome of birds is often thought to evolve via stronger drift (Mank et al. 2009; Hayes et al. 2020; Chase et al. 2023) but the Z of Lepidoptera is sometimes found to be adaptively evolving (Sackton et al. 2014; Mongue et al. 2022) and sometimes not (Rousselle et al. 2016; Pinharanda et al. 2019). Within these patterns are significant disagreements though, as there have been arguments for an adaptive fast Z in birds (Dean et al. 2015) as well as for a lack of a fast Z at all in Lepidoptera (Rousselle et al. 2016). Both of these clades have single-origin Z chromosomes (Griffiths et al. 1998; Fraïsse et al. 2017), so, discounting methodological variation, differences between studies may be attributable to differences in choices of study species and their biology. Below, we unpack our novel finding that the *Lepeophtheirus* faster Z is not driven by increased adaptation on the sex chromosome and relate this finding to established sex chromosome results. To understand why the salmon louse Z is not more adaptively evolving, we consider patterns of evolution of each of the classes of sex-biased genes and how they relate to organismal biology.

First, male-biased genes should preferentially accumulate on the Z chromosome because of its tighter association with males compared to females, a pattern observed in numerous species (Storchová and Divina 2006; Ellegren 2011; Mongue and Walters 2017) including *L. salmonis* in this study. The Z is diploid in males though, so any male-biased Z-linked gene should be subject to the same selective forces as their autosomal counterparts, albeit with the potential decrease in the efficacy of selection do to a lower effective population size of Z chromosomes compared to autosomes (Vicoso and Charlesworth 2009). In other words, male-biased genes are not traditionally expected or observed to contribute to faster Z adaptation. Mongue et al. (2022) found some evidence for increased male-biased adaptation on the Z of the monarch butterfly, *Danaus plexippus*, but this species is known to have extremely high rates of polyandry, with some females mating up to 14 times in quick succession (Hill Jr. et al. 1976; Smith 1984). Such extreme sperm competition and/or cryptic choice may increase positive selection on male-biased traits in this species (Mongue et al. 2019). In general, however, low to intermediate levels of polyandry, as have been observed in *L. salmonis* (Todd et al. 2005), are more likely to merely marginally decrease the strength of drift and permit fewer weakly deleterious variants in the population (Dapper and Wade 2016). This prediction lines up well with the decreased *pN*/*pS* of male-biased genes on the Z and autosomes as well as the marginally higher α value seen for autosomal male-biased genes compared to their unbiased counterparts.

Female-biased genes will be expressed in a haploid state on the Z and thus should experience more efficient haploid selection (Charlesworth et al. 1987). Conversely, the Z chromosome is predicted to have a lower effective population size than the autosomes, so genes on it should experience stronger drift (Vicoso and Charlesworth 2009) which could somewhat counteract the benefits of haploid selection. Many of the studies of avian Z chromosomes do not parse evolutionary patterns by sex-bias class, obscuring this dynamic. In Lepidoptera though, evidence is more readily presented but no clearer. In two lineages, female-biased Z-linked genes are no more adaptively evolving than their autosomal counterparts (Pinharanda et al. 2019; the Carolina sphinx moth Z in Mongue et al. 2022). In two other cases, female-biased genes *do* evolve more adaptively on the Z (Sackton et al. 2014; the monarch neo-Z Mongue et al. 2022). Part of this discrepancy could be explained by the underpowered nature of tests for selection combined with low counts of female-biased genes on the Z. Variability between genes swamps any aggregate signal of positive selection. This scenario is certainly plausible for *L. salmonis*, which does not have many Z-linked female-biased genes to analyze.

On the biological side of explanations, it is worth noting that some of the best evidence for faster Z adaptation of haploid-expressed genes comes from the neo-Z chromosome of monarch butterflies (Mongue et al. 2022). The genus containing monarchs has a previously autosomal segment of chromosome fused to the ancestral Z chromosome and termed the neo-Z (Mongue et al. 2017). This portion of the chromosome is not only less male-biased than the ancestral Z, but also regulated in a different manner. Because the Z chromosome varies in ploidy between the sexes, regulatory mechanisms have evolved normalize expression between sexes, often termed dosage compensation; in Lepidoptera, the Z chromosomes are typically down-regulated in males to match the lower expression of the single copy Z in females (Gu and Walters 2017). The neo-Z however is upregulated in females to match autosomal expression (Gu et al. 2019). The strength of selection is thought to relate to the level of expression, with more highly expressed genes being more exposed to selection (Vicoso and Charlesworth 2009). Thus, it stands to reason that an upregulated Z chromosome should be more visible to selection than a down-regulated one. In this study we found a pattern of down-regulation of the Z in *L. salmonis*, suggesting that there may be less selective scrutiny of haploid-expressed female-biased genes on the Z of salmon lice.

Moreover, sex-biased genes should be particularly responsive to variation in reproductive success of the sexes. Males are typically thought to have a higher variance in reproductive success than females (Bateman 1948), which could drive the effective population size of the Z even lower than 3/4^th^ that of the autosomes. Yet our nucleotide-diversity-based estimates of relative effective population size of the Z and autosomes do not clearly show this pattern. Rather unusually, the Z has only a marginally lower effective population size compared to the autosomes (0.95), suggesting that the Z is more often passed on in a diploid than haploid state, i.e. females have significantly higher variance in reproductive success compared to males. This counterintuitive conclusion is backed up by organismal study as well. A long-term study of wild *L. salmonis* concluded that “at least 30% of reproductive female sea lice experience mate limitation” thanks to their parasitic lifestyle hindering dispersal to find mates (Cox et al. 2017). As such, any beneficial Z-linked alleles not directly related to mate acquisition are likely at a high risk of being lost to genetic drift, haploid selection or no. This prediction is also supported by the observation that female-biased genes, both on the Z and autosomes are more permissive of (likely neutral or deleterious) non-synonymous polymorphisms than male-biased or unbiased genes.

Finally, unbiased genes experience both diploid male selection and haploid female selection on the Z, depending on the sex in which they are expressed. As such, their evolutionary patterns should be somewhere in between male- and female-biased genes. Which of the two sexes they more closely resemble is an unresolved question though. Again, evidence comes mainly from Lepidoptera rather than Aves. In Lepidoptera with faster Z adaptation, unbiased genes were a contributing factor (i.e. evolved more adaptively on the Z than autosomes) in two of the three lineages, but note that the monarch ancestral Z is driven by male-biased genes (Sackton et al. 2014; Mongue et al. 2022), so unbiased gene adaptation does not appear to be a strict requirement for faster Z adaptation.

Another line of thinking runs orthogonal to the consideration of sex-biased genes and sexual conflict on the sex chromosomes. Because unbiased genes on the sex chromosomes encode traits expressed in both sexes that also happen to benefit from haploid selection in females, they have been proposed to contribute disproportionately to local adaptation (Connallon et al. 2018). Indeed in the Carolina sphinx moth unbiased Z-linked genes are known to be more adaptively evolving (Mongue et al. 2022) and multi-population comparisons have identified a population-specific, high frequency Z-linked inversion that is enriched for unbiased genes relative to the rest of the Z chromosome (Mongue and Kawahara 2022). From this perspective, it is perhaps unsurprising that we fail to see a similar effect in *Lepeophtheirus*. With such a high rate of female reproductive failure, many beneficial locally adapted alleles can be lost when females fail to mate.

## Conclusion

Our analysis of the salmon louse adds a new independent datapoint for understanding sex chromosome evolution. By parsing genes by sex-bias of expression we are able to thoroughly explore theoretical predictions and by intersecting these molecular results with behavioral ecology, we uncover the role of mating system dynamics in sex chromosome evolution. In summary, the *Lepeophtheirus* faster-Z appears to be driven mainly by drift, with no evidence for a role of haploid expression in females causing faster adaptive evolution. Though haploid selection should be a powerful force for both purifying and positive selection, the relative paucity of Z-linked female-biased genes, lack of dosage compensation, and high rate of reproductive failure of females likely counteracts these benefits via strong drift. More generally, we hope this study highlights how the diversity of arthropod sex determination can be used to advance population genetic theory through comparative genomics and how an increasing wealth of whole-genome sequencing data generated for distinct purposes can be synthesized to answer new questions.

## Acknowledgements

The authors thank University of Florida Research Computing for providing computational resources and support that have contributed to the research results reported in this publication. URL: http://www.rc.ufl.edu. We would also like to thank the Florida Museum Marine Invertebrate Collection and Gustav Paulay in particular for providing expertise and access to outgroup sea louse samples. Finally, we wish to thank the lab of Laura Ross and the Ashworth computing cluster group at the University of Edinburgh for providing a supportive environment in which to explore evolutionary genetics.

## Data availability

All data used in analyses are publicly available via the accessions in Table 1. The *Lepeoptheirus parviventris* Illumina WGS reads generated for this study were ultimately not used as they were too diverged from *L. salmonis* but they may be a useful tool for others and will be made available upon publication.

